# Intravital Depletion of Cholesterol Reduces Membrane Dynamics and Increases Mechanosensitivity in Osteocytes In Vivo

**DOI:** 10.64898/2026.07.21.739862

**Authors:** Melia D. Matthews, Katelyn Lunny, Lakhi Raju, Nada Naguib, Saniya Tariq, Ulirich B. Wiesner, Karl J. Lewis

## Abstract

Osteocytes detect mechanical forces within bone through signaling processes organized at the plasma membrane. Membrane cholesterol regulates membrane organization, dynamics, and mechanical properties, yet its role in osteocyte mechanotransduction *in vivo* remains unknown. Here, we developed an intravital multiphoton imaging approach to quantify membrane-associated uptake, retention, and clearance alongside load-induced Ca^2+^ signaling in osteocytes within the intact metatarsal. Using fluorescent nanoparticles and a membrane-labeling probe, we tracked these processes before and after pharmacological cholesterol depletion. Cholesterol depletion reduced osteocyte membrane uptake and retention and altered clearance in both sexes, while affecting load-induced Ca^2+^ signaling in a sex-dependent manner. In females, cholesterol depletion increased both the proportion of osteocytes responding to mechanical loading and the magnitude of their responses, whereas neither outcome changed in males. These findings identify plasma membrane cholesterol as a regulator of osteocyte membrane dynamics and mechanical responsiveness *in vivo*. More broadly, this work establishes an approach for directly examining how membrane composition and turnover regulate mechanotransduction in cells embedded within their native tissue environment.

## Introduction

Bone is a dynamic tissue that continuously adapts to mechanical inputs from its local environment(1, 2). Tissue-level strain generates pressure gradients that drive fluid flow through the lacuno-canalicular system (LCS)(3, 4). Osteocytes, which reside within the LCS, function as the primary mechanosensors of bone by detecting these forces, regulating osteoblast and osteoclast activity, and coordinating skeletal adaptation(5).

Fluid flow-induced shear stress within the LCS is amplified at integrin attachment sites along osteocyte dendrites, allowing small tissue deformations to activate mechanosensitive signaling pathways(6–8). Consistent with computational and mathematical models, application of piconewton-scale forces to osteocyte dendrites, but not the cell body, triggers intracellular Ca^2+^ signaling waves that propagate throughout the cell(7, 9). These findings highlight the importance of dendrite force amplification in osteocyte mechanotransduction. However, despite substantial progress in defining how osteocytes sense force, key questions remain in the field regarding the cellular mechanisms that regulate osteocyte mechanobiology.

One potential regulator is the plasma membrane. Beyond serving as a passive mechanical interface, the plasma membrane is a dynamic structure whose composition and organization influence transmembrane protein trafficking, mechanosensitive signaling, and cellular force transduction(10–13). In fibroblasts, endothelial cells, and other mechanosensitive cell types, membrane properties such as fluidity, tension, and lipid organization modulate downstream mechanosensitive responses(14, 15). Cholesterol is a key determinant of these membrane properties, contributing to lipid raft formation, membrane organization, and endocytic activity in concert with proteins like Caveolin-1(16–18). Importantly, membrane cholesterol levels have been shown to regulate cellular mechanosensitivity by altering membrane tension and transmembrane protein organization(15). Experimental depletion of cholesterol with cyclodextrins increases membrane fluidity, disrupts lipid raft organization, and alters membrane tension, making it a widely used approach for probing the role of membrane dynamics in cell function and mechanosensitivity(19–22)

Despite these insights from broader cell biology, the regulatory role of the plasma membrane in osteocytes remains unclear. Most studies of osteocyte membranes have focused on passive membrane deformation during fluid shear stress, predominantly using computational stimulations derived from 3D confocal imaging of osteocytes within the LCS(8, 23–25). In contrast, the active dynamics of the plasma membrane have yet to be fully explored.

While osteocyte sensitivity to mechanical stimulation under different conditions has been well studied*in vitro*(26, 27) and*ex vivo*(28–30), direct observation of real-time osteocyte dynamic activity*in vivo* has only recently become feasible. Advances in multiphoton microscopy and surgical procedures have allowed observation of osteocyte Ca^2+^ signaling under mechanical load in their native 3D environment, preserving vasculature and endocrine signals(30, 31). However, real-time visualization of membrane dynamics, cholesterol depletion, and their connection to osteocyte mechanosensitivity have yet to be described*in vivo*.

To address this gap, we developed an intravital imaging methodology to directly observe and quantify the interplay between cholesterol-dependent membrane dynamics and osteocyte mechanosensitivity. We previously established a platform using locally injected fluorescent nanoparticles to study osteocyte activity*in vivo*(32). These nanoparticles were rapidly taken up and cleared by osteocytes, providing a functional read-out of membrane activity and trafficking. We further observed that nanoparticle dynamics could be modulated through locally pharmacological perturbation, including a cholesterol depletion(33).

Here, we combined this intravital imaging platform with the membrane dye FM1-43 to investigate the relationship between membrane dynamics and mechanosensitivity in metatarsal osteocytes and to determine their dependence on plasma membrane cholesterol. We tested two complementary hypotheses: first, that cholesterol depletion disrupts osteocyte membrane dynamics and trafficking; and second, that cholesterol depletion results in increased osteocyte sensitivity to mechanical stimulation. Our findings reveal a strong coupling between membrane activity and mechanosensitivity in osteocytes*in vivo*. Specifically, cholesterol depletion both reduced osteocyte membrane dynamics and modulated their load-induced Ca^2+^ signaling in a sexually dimorphic manner.

## Results

### Subcutaneous application of Methyl-β-cyclodextrin reduces plasma membrane cholesterol in cortical bone osteocytes

To develop an*in vivo* model for perturbing osteocyte membrane dynamics and trafficking, skeletally mature male and female mice received subcutaneous treatment in the hind paw with the cholesterol-depleting agent methylβ-cyclodextrin (MβCD) or DPBS control. Following dissection, fixation, and cryosectioning of the third metatarsal (MT3), cortical bone osteocyte cholesterol levels were assessed by Filipin III staining.

As expected from previous reporting in other tissues, an*in vivo* subcutaneous injection of M*β*CD (30mM, 15µL) reduced Filipin III signal intensity in MT3 cortical bone osteocytes (Figure 1). No changes were observed in fluorescent GCaMP6f Ca^2+^ signal with treatment (Figure 1G,N). The M*β*CD-induced reduction in fluorescent signal was observed across both male and female mice. Notably, the decrease in Filipin III fluorescent staining was stronger in females, with a 95% decrease in cholesterol compared to a 65% reduction in males.

**Fig. 1.**
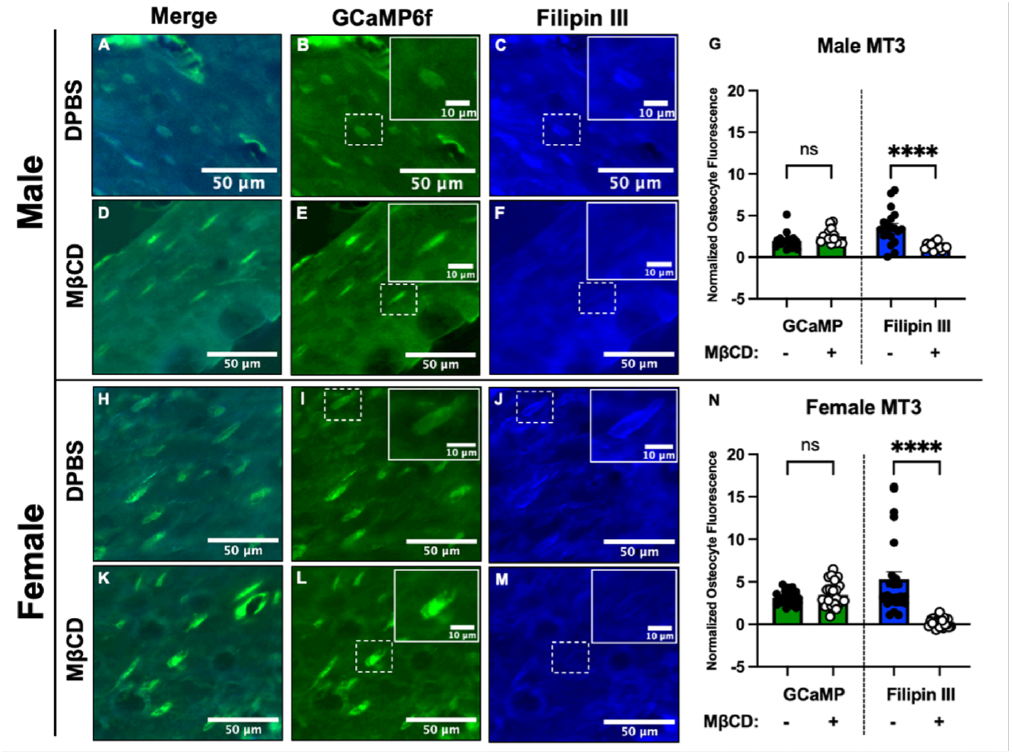
A subcutaneous injection of M*β*CD decreases MT3 osteocyte cholesterol levels in vivo. Cryosectioned MT3 cortical bone from a male GCaMP6f mouse following a 30-minute exposure to control 1 *×* DPBS (A–C) or 30 mM M*β*CD (D–F). Osteocytes were identified by green fluorescent GCaMP6f Ca^2+^ signal. Cellular cholesterol was fluorescently stained with Filipin III. Scale bars = 50 *µ*m. Representative GCaMP6f- and Filipin III-stained cells identified by dashed white boxes are enlarged in the upper-right corners; inset scale bars = 10 *µ*m. Normalized fluorescent signals from male osteocyte ROIs were quantified for GCaMP6f and Filipin III and compared between treatment groups using a one-way ANOVA (G; \*\*\*\**p <* 0.0001). Staining was repeated in a female MT3 following exposure to 1*×* DPBS (H–J) or 30 mM M*β*CD (K–M). Scale bars = 50 *µ*m. Representative GCaMP6f- and Filipin III-stained cells identified by dashed white boxes are enlarged in the upper-right corners; inset scale bars = 10 *µ*m. Normalized signals from female osteocyte ROIs were quantified for GCaMP6f and Filipin III and compared between treatment groups using a one-way ANOVA (\*\*\*\**p <* 0.0001).

### Cholesterol depletion depresses membrane and nanoparticle dynamics in osteocytes*in vivo*

To assess the impact of cholesterol depletion on membrane trafficking in osteocytes*in vivo*, we created a novel timelapse imaging protocol using subcutaneously injected fluorescent dyes. These experiments utilized the membrane-dye FM1-43 with either integrin-targeted RGD-C Dots or cytosol-targeted TATC ′ Dots. These C’Dot types were chosen to target both regulated trafficking (RGD) and endosomal evasion (TAT) under the experimental state of cholesterol depletion. A short, 5 minute incubation period before MT3 isolation and imaging was performed to observe rapid internalization and trafficking of the membrane and nanoparticles (Figure 2A). Repeated imaging of the fluorescent signals was conducted over the time span of an hour forM*β*CD treated and naïve mice (Figure 2B).

**Fig. 2.**
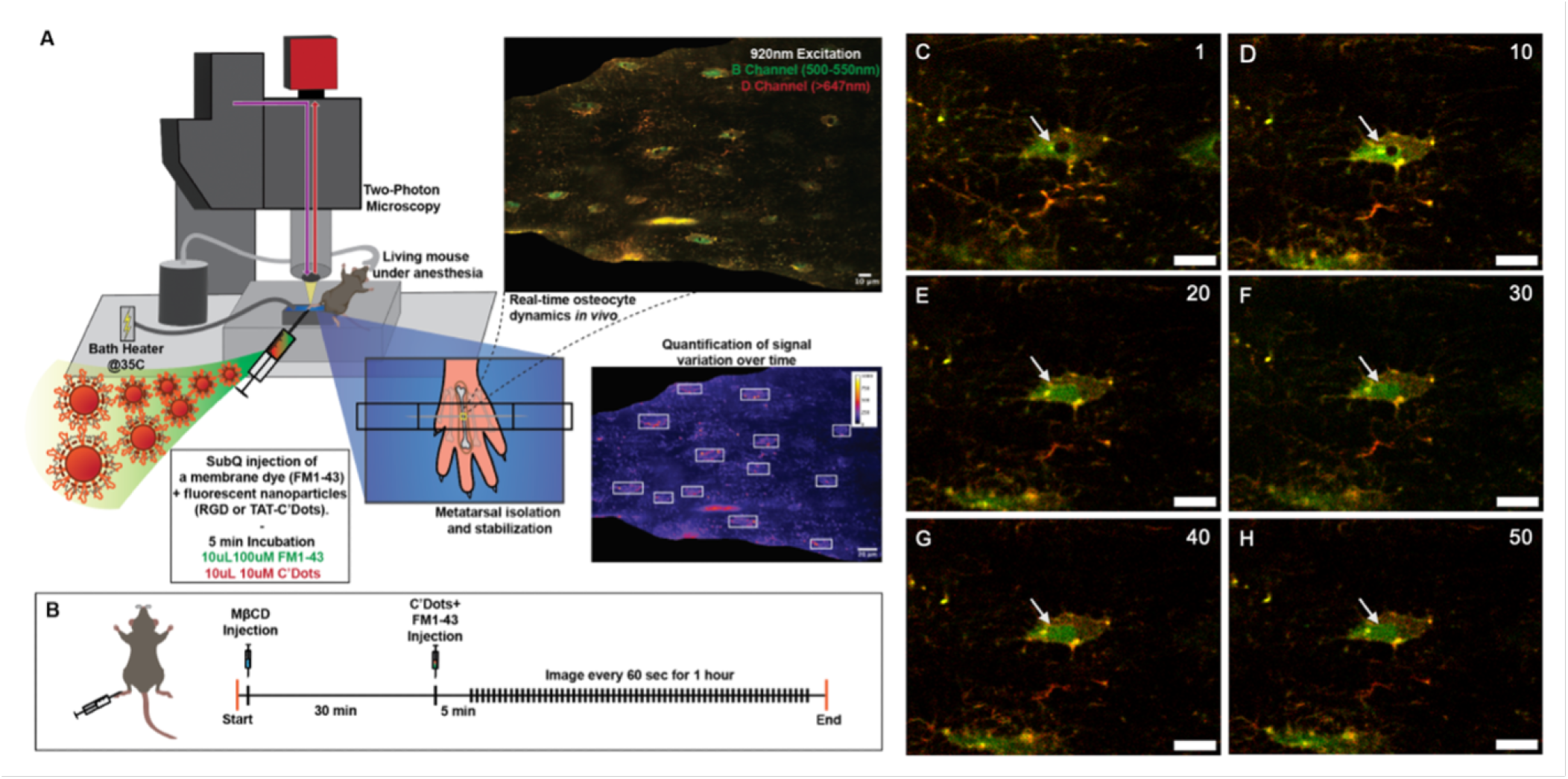
Timelapse imaging of membrane and nanoparticle dynamics in osteocytes in vivo using multiphoton microscopy. Fluorescent signal from subcutaneously injected dyes and nanoparticles can be visualized in the cortical bone of the MT3 using a previously described surgery and stabilization method (A)(31, 32). A custom immersion heater was built to maintain the DPBS bath temperature at 35^*°*^C and promote membrane trafficking activity. Schematic of the experimental timeline showing M*β*CD injection, fluorescent marker injection, and time-lapse imaging (B). Representative images acquired every 10 min with overlaid red and green channels throughout the experiment (C–H). The white arrow indicates the initial location of a punctate signal that moves over the course of the experiment. Scale bar = 10 *µ*m.

Both green (membrane marker) and red (nanoparticle) fluorescent signals were collected simultaneously and tracked in individual osteocytes at the micron and submicron scale (Supplemental Figure 1). Movement of membrane and nanoparticle fluorescent signal was observed in cells over the course of the time lapse imaging, confirming our ability to see real-time dynamic activity in osteocytes*in vivo* (Figure 2C-H, Supplemental Videos 1,2).

Fluorescent signal quantified in individual cells revealed that M*β*CD-treated mice exhibited less change in signal intensity than control mice, suggesting reduced membrane trafficking activity. M*β*CD-treated groups displayed shallower slopes in linear regressions of mean cellular fluorescence intensity over the course of the experiment (Figure 3A–H). Cholesterol depletion also reduced the initial uptake of fluorescent signal. This trend was consistent across sexes for both membrane and nanoparticle signals. However, in females, membrane and C′ Dot signal intensities in the control and M*β*CD-treated TAT groups converged during the second half of the experiment (Figure 3D,H), suggesting that the effects of cholesterol depletion diminished over time. This trend was not observed in males or in the RGD-C′ Dot groups.

**Fig. 3.**
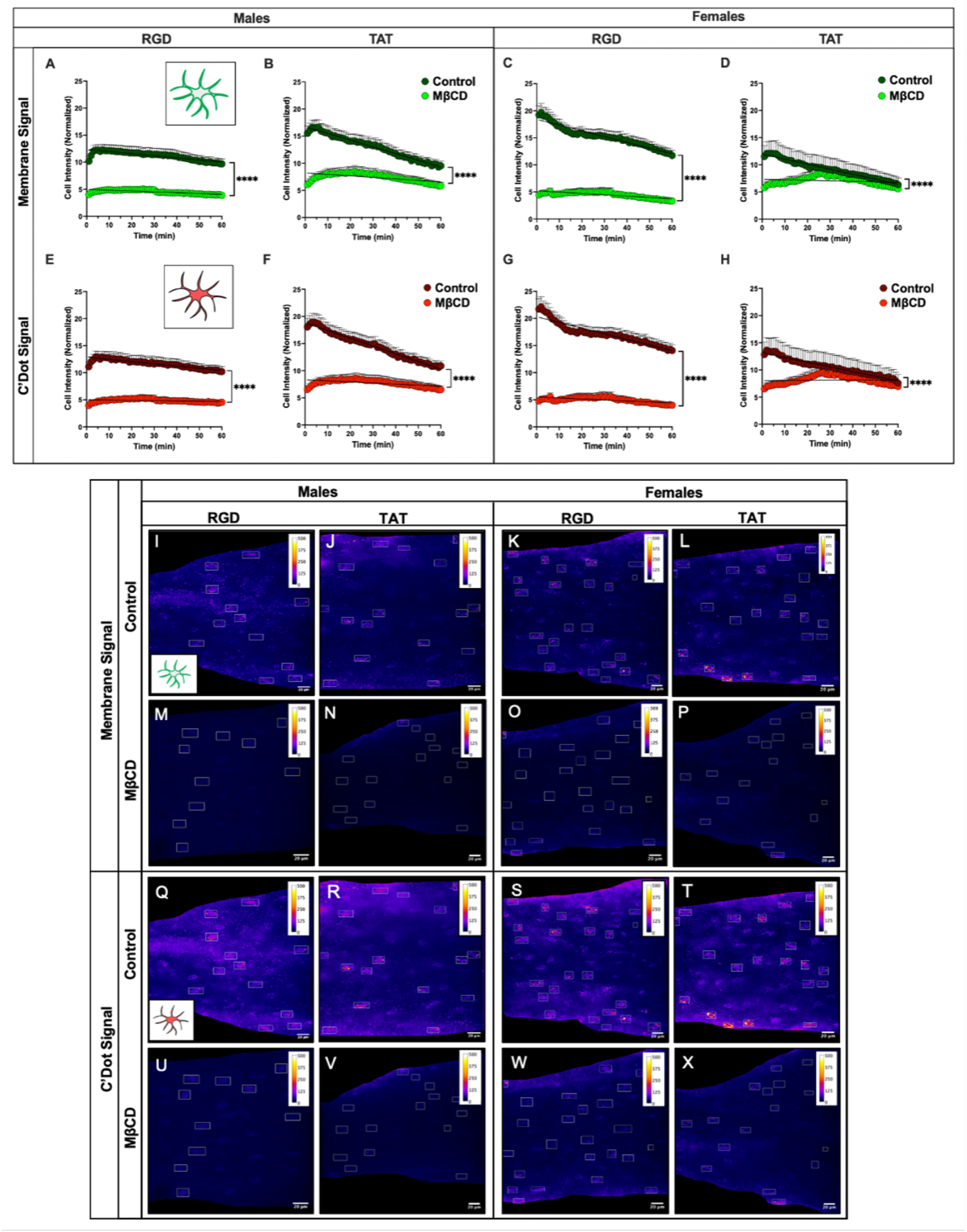
Cholesterol depletion with M*β*CD reduces membrane and nanoparticle signal uptake and dynamics in osteocytes *in vivo*. Changes in fluorescence intensity over time were quantified by linear regression of mean cellular fluorescence for membrane signal (A–D) and C′ Dot signal (E–H). Asterisks indicate significant differences in the slopes of linear regressions between control and M*β*CD-treated groups. *n* = 3 mice per group, with 10–30 cells analyzed per mouse. Individual data points represent measurements from single cells. Error bars represent SEM. \**p <* 0.05, \*\**p <* 0.01, \*\*\**p <* 0.001, \*\*\*\**p <* 0.0001. Standard deviation *z*-projections of the MT3 ROI were converted to heatmaps to visualize cellular dynamics during membrane and nanoparticle time-course imaging. Calibration bars indicate mean gray values ranging from 0 to 500. Representative heatmaps of membrane signal in control (I–L) and M*β*CD-treated (M–P) groups. Representative heatmaps of ′ Dot signal in control (Q–T) and M*β*CD-treated (U–X) groups. Cells selected for quantification are indicated by white boxes. Scale bars = 20 *µ*m.

Looking specifically at the first imaging time point, M*β*CD-treated groups exhibited lower initial signal uptake than control groups across both sexes and C ′ Dots. These C′ Dot types were chosen to target both regulated trafficking (RGD) and endosomal evasion (TAT) un Dot functionalizations (Supplemental Figure 2A). These findings suggest that cholesterol depletion reduces the ability of osteocytes to rapidly internalize nanoparticles. By the 60 min time point, all male membrane and C ′ Dots. These C ′ Dots. These C′ Dot types were chosen to target both regulated trafficking (RGD) and endosomal evasion (TAT) un-Dot types were chosen to target both regulated trafficking (RGD) and endosomal evasion (TAT) un-

Dot groups retained higher fluorescence intensity than their corresponding M*β*CD-treated groups (Supplemental Figure 2B), indicating that the effects of cholesterol depletion persisted throughout the experiment. In females, however, the TAT-C′ Dot group no longer differed from its M*β*CD-treated counterpart in either the membrane or C Dot channel, consistent with the convergence observed in the fluorescence intensity time-course plots (Figure 3D,H). The equivalent signal intensity at the 60 min time point suggests that the effects of cholesterol depletion diminished over time in the female TAT-C′ Dot group.

Membrane dynamics and trafficking variability in osteocytes over the hour-long experiment were also assessed using standard deviation *z*-projection heatmaps of cellular fluorescence intensity, in which brighter colors indicate greater changes in signal over time (Figure 3I–X). Cholesterol depletion reduced dynamic fluorescence across all groups, resulting in darker heatmaps for both membrane (Figure 3M–P) and C Dot (Figure 3Q–X) signals in the M*β*CD-treated groups. This trend was consistent across both sexes and C Dot functionalizations.

Quantification of these heatmaps confirmed that M*β*CD reduces overall signal variation and membrane trafficking in osteocytes *in vivo*. Cholesterol depletion decreased variation in fluorescence intensity for both membrane and C′ Dot signals in male and female mice (Figure 4A–B). These findings are consistent with the effects of M*β*CD on fluorescence intensity observed in Figure 3, supporting the use of signal variation heatmaps as a quantitative measure of membrane dynamics. Interestingly, C′ Dot functionalization did not affect signal variation (Figure 4C–D), suggesting that the short incubation period may not have been sufficient for nanoparticle targeting to produce distinct trafficking patterns. Moderate sex differences in signal variability were observed; female mice injected with RGD-C′ Dots exhibited greater variation in both membrane and nanoparticle signals than males (Supplemental Figure 3). In contrast, no sex differences in signal variation were detected in the TAT-C′ Dot or M*β*CD-treated groups.

**Fig. 4.**
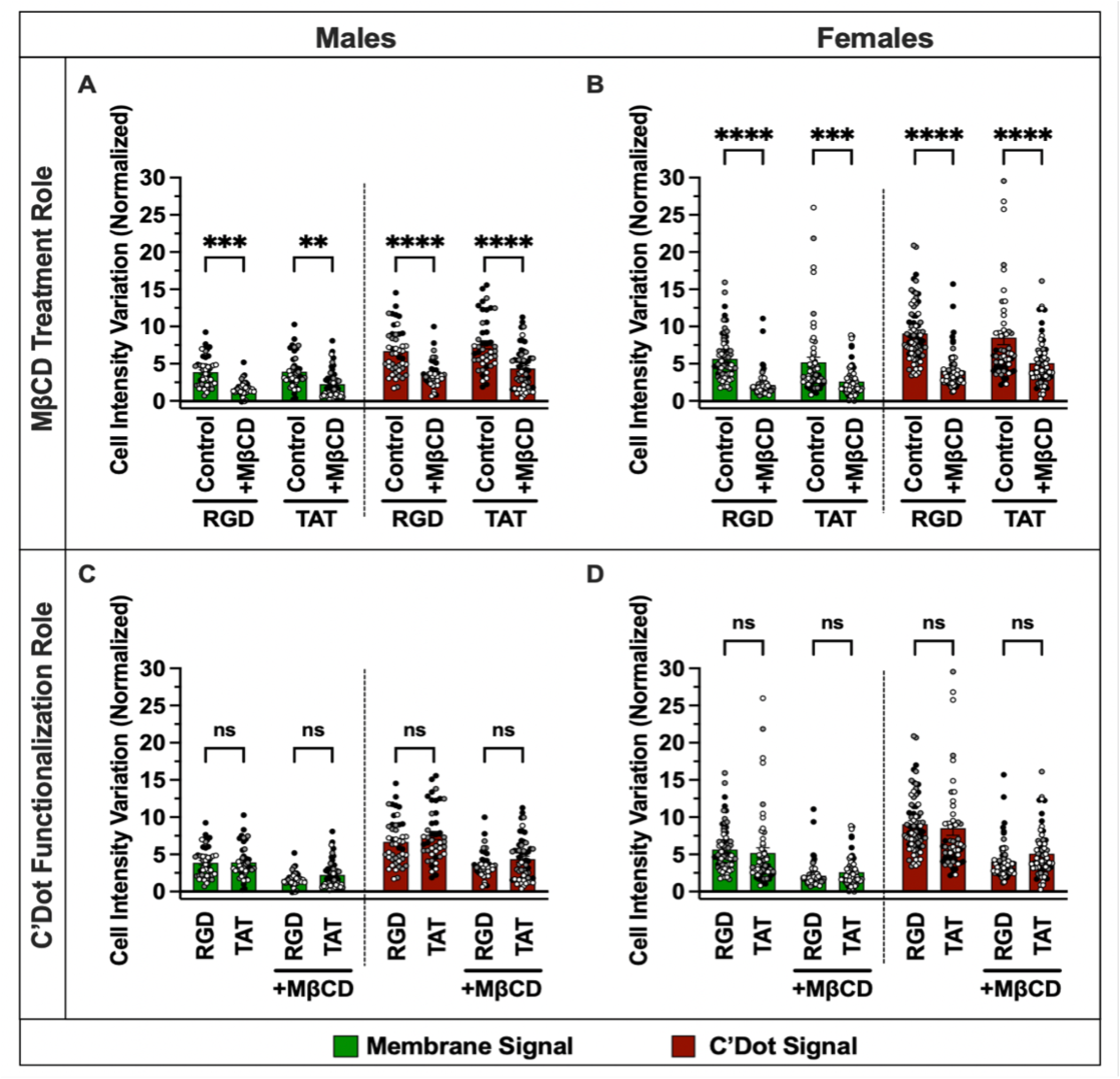
Cholesterol depletion, but not C^2+^ Dot functionalization, alters signal intensity variation in osteocytes over the course of the time-lapse experiment. Signal intensity variation was quantified as the standard deviation *z*-projection of cellular ROIs over the experimental time course for both membrane and C′ Dot signals. The effects of M*β*CD treatment (A–B) and C′ Dot functionalization (C–D) on signal intensity variation were assessed using one-way ANOVAs with multiple comparisons. *n* = 3 mice per group, with 10–30 cells analyzed per mouse. Individual data points represent measurements from single cells, with colors indicating individual mice. Error bars represent SEM. \**p <* 0.05, \*\**p <* 0.01, \*\*\**p <* 0.001, \*\*\*\**p <* 0.0001.

**Fig. 5.**
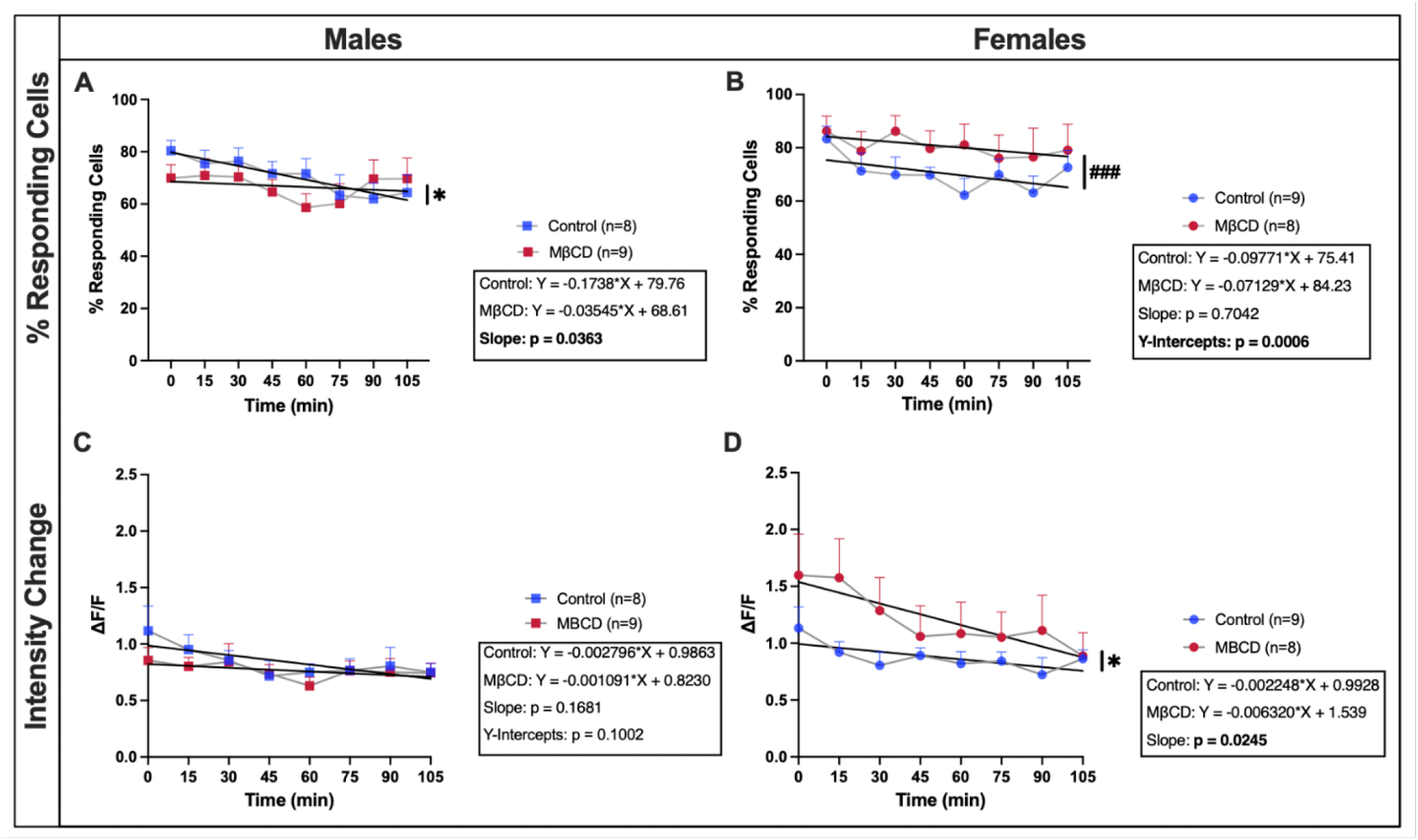
Application of M*β*CD alters osteocyte Ca^2+^ signaling and mechanosensitivity *in vivo*. The percentage of responding osteocytes during each loading session in males (A) and females (B). Load-induced changes in Ca^2+^ fluorescence (Δ*F/F* ) between non-loaded and loaded imaging time points in males (C) and females (D). Data were analyzed using linear regression of the mean response at each time point, with comparisons between control and M*β*CD-treated groups. Asterisks indicate significant differences in slope, whereas pound symbols (#) indicate significant differences in *y*-intercept. *N* = 8–9 mice per group. Error bars represent SEM. \**p <* 0.05, \*\**p <* 0.01, \*\*\**p <* 0.001, \*\*\*\**p <* 0.0001; #*p <* 0.05, ##*p <* 0.01, ###*p <* 0.001, ####*p <* 0.0001.

### Cholesterol depletion increases osteocyte mechanosensitivity in female mice, with lesser affect in males

To measure the direct impact of cholesterol depletion and reduced membrane dynamics on osteocyte mechanosensitivity, we used*in vivo* loading with simultaneous multiphoton observation of Ca^2+^ signaling. Physiological loading was applied to GCaMP6f mouse metatarsal bones*in vivo* after a local subcutaneous injection of M*β*CD or a vehicle control. Fluorescent Ca^2+^ signaling in osteocytes was imaged before and during repeated loading bouts and the percentage of osteocytes responsive to loading during each bout was calculated.

Application of M*β*CD altered the percentage of osteocytes that responded to physiological loading over time in both sexes. While the control group in males had a steady decrease in the percentage of responding cells over the course of the experiment, the trend line was nearly flattened in M*β*CD treated males (Figure 4A). There was no difference in osteocyte Ca^2+^ fluorescent intensity change in drug treated male mice compared to control mice during loading (Figure 4C). In females however, the impact of cholesterol depletion was more distinct; while the slopes remained the same, M*β*CD treated mice had a higher percentage of responsive osteocytes over the entire course of the experiment, as measured by the y-intercept of linear regressions (Figure 4B). M*β*CD application resulted in up to 33% more osteocytes responding to the same level of physiological loading in females. Additionally, osteocyte Ca^2+^ fluorescence intensity change was higher in M*β*CD treated females, although the intensity change decreased over the course of the experiment relative to control treated mice (Figure 4D).

## Discussion

In this study, we report the first real-time observations of osteocyte membrane dynamics and trafficking*in vivo*, revealing a strong relationship between plasma membrane cholesterol, membrane dynamics, and osteocyte mechanosensitivity. Building upon earlier work establishing the uptake kinetics of C’Dot nanoparticles, these findings advance the ability to interrogate osteocyte activity within their native 3D environment while preserving physiological paracrine and endocrine signaling. This work further validates subcutaneous drug delivery as a robust approach for localized,*in vitro*-style cell modulation*in vivo*. In addition, we demonstrated that subcutaneously delivered fluorescent dyes such as FM1-43 can be used to directly monitor osteocyte membrane dynamics in living bone. Together, these findings establish fluorescent nanoparticles and membrane probes as powerful tools for studying osteocyte activity*in vivo* and support a model in which cholesterol-dependent membrane dynamics and mechanosensitivity are tightly coupled within osteocytes.

Fluorescent Filipin III staining confirmed the cholesterol depleting role of M*β*CD in our system, aligning with many previously published studies using cyclodextrins to reduce cellular cholesterol levels(19, 34–36). Osteocytes in both male and female metatarsal bone displayed reduced levels of cholesterol after treatment; this aligns with our previously reported results that subcutaneous application of M*β*CD influenced osteocyte membrane dynamics*in vivo* across both sexes through increased nanoparticle uptake and retention(33). These confirmatory results supported further investigation of cholesterol depletion on osteocyte activity with our M*β*CD injection and incubation protocol.

Using our novel timelapse imaging methodology to visualize membrane dynamics in real time in osteocytes*in vivo*, we found that cholesterol depletion with M*β*CD treatment dramatically reduced the uptake and variability of membrane and C’Dot signal. These results align with the widely studied impact of M*β*CD, which disrupts lipid rafts and reduces overall membrane activity and endocytosis(34, 35, 37, 38). M*β*CD is known to perturb the formation of caveolae invaginations and clathrin coated pits, reducing presence of endocytic vesicles(39–41). While clear differences were observed in membrane dynamics after M*β*CD treatment as expected, fewer differences were observed between nanoparticle functionalizations. The short 5-minute incubation period, selected to maximize uptake activity and FM1-43 membrane fluorescence, may have limited the divergence between integrin-targeted (RGD) and membrane-penetrating (TAT) particles relative to longer incubation conditions. Prior short-term clearance data from our lab support this interpretation(33). The reason that control TAT-C′ Dot and membrane signal converged with M*β*CD-treated signal in the second half of the timelapse experiment in females, while RGD-C′ Dot and membrane signals remained distinct, is unclear (Figure 33D,H).

Cholesterol depletion also impacted osteocyte mechanosensitivity to physiological load. Our results were sexually dimorphic, with M*β*CD-treated females displaying an increased percentage of osteocytes responding to repeated bouts of 1000µε compared to controls while male groups had a less clear distinction. Notably, a previous experiment performed by Xing et al., found that cultured osteoblast-like MC3T3-E1 cells displayed a strong decrease in mechanosensitivity after treatment with M*β*CD(42). They found that Ca^2+^ peaks in response to oscillatory fluid flow nearly disappeared after cholesterol depletion*in vitro*. We observed the opposite effect (in females), suggesting that cell type and native 3D environment play a critical role in studying the relationship between cell membrane and mechanosensitivity. Notably, our cholesterol depletion results with Filipin III staining suggested that females have a stronger reduction in cholesterol compared to males after M*β*CD treatment (Figure 1). This differential response may influence the subsequent sexual variability we observed in osteocyte response to loading. In other cell types, M*β*CD is known to increase sensitivity to mechanical stimulation. Biswas et al., link increased membrane tension from cholesterol depletion to higher sensitivity, observing greater rates of cell rupture during hypo-osmotic shock(19). Other groups have shown that MβCD-increased membrane tension augmented TRPV4 stretch-activated ion channel sensitivity and expression in human trabecular meshwork cells(43).

We hypothesize that cholesterol depletion may increase membrane tension and disrupt the trafficking and recycling of transmembrane proteins like integrins, allowing osteocyte dendrites to maintain higher levels of sensitivity to loading compared to their steady state. Integrin attachment sites occur at regular intervals along the dendrites, colocalizing with other mechanosensitive proteins including CaV3.2, P2X7R, and Pannexin1 that contribute to the osteocyte mechanosome(44, 45). Proper function of these proteins depends on their organization and mobility, requiring regular internalization and recycling to control expression and localization(46–48). However, the interior of osteocyte dendrites (∼100nm diameter) is almost entirely filled by actin cytoskeletal filaments that provide structural support to the cellular processes(44, 49–51). This spatial constraint leaves no room for the canonical vesicular trafficking and recycling of integrins and other transmembrane proteins(45). Trafficking vesicles are on the order of ∼30-50nm in diameter, simply too large to fit along with the cytoskeletal backbone of osteocyte processes(52, 53). These constraints suggest that mechanisms beyond conventional vesicular trafficking may be required to regulate protein localization along osteocyte dendrites, implicating the importance of plasma membrane organization and dynamics in this process.

The fluid mosaic model may help explain how transmembrane proteins are trafficked along osteocyte dendrites, with proteins transported along the cell surface rather than inside bulky vesicles(11, 15). Supporting this idea, myosin expression has been observed within osteocyte dendrites, suggesting active membrane-associated transport without internalization could be feasible(54). Modulating membrane composition and fluidity may therefore represent a strategy to restore dysregulated mechanosome protein localization and mechanosensitivity in disease states such as OVX and aging(31).

Previous intravital imaging studies identified sexual dimorphism in osteocyte endocytic pathways between male and female mice(33). Consistent with these findings, females in the present study displayed greater membrane dynamics and trafficking activity than males during timelapse imaging, including increased signal variability in both RGD-C Dots and membrane markers (Figure 3K-L). Females also exhibited higher qualitative membrane trafficking activity (data not shown). These findings align with prior reports of elevated osteocyte activity in females during perilacunar remodeling associated with pregnancy and lactation(55–57), suggesting that female osteocytes may maintain intrinsically higher trafficking activity more broadly. The mechanisms underlying this sexual dimorphism remain unclear but may involve gonadal hormones or differential gene expression.

There were several limitations of the experimental work that should be highlighted. The timelapse studies are comprised of a relatively small sample size (n=3 per group), although we did quantify activity and signal in at least 10 cells per mouse. Additionally, in the loading and Ca^2+^ signaling experiment, our 3-point bending was limited to a single magnitude, repeated over time in order to observe any effect of drug washout. In previous work, we have seen that the impact of an intervention on the percentage of responding osteocytes can change at different loading magnitudes(31, 58). Expanding the range of tested loading magnitudes is an important future experiment to understand the holistic impact of cholesterol depletion on osteocyte mechanosensitivity*in vivo*.

In total, the findings presented here expand a paradigm of interaction between osteocyte membrane dynamics and mechanosensitivity. Cholesterol depletion reduces membrane dynamism and alters mechanosensitivity in osteocytes, specifically increasing the percentage of responding cells in females. We continue to observe robust sex differences in membrane dynamics, the origins and mechanisms of which should be further investigated. These studies build on our previous work and solidify this platform as a robust tool for visualizing osteocyte subcellular activity*in vivo* under both loading and drugged conditions. Further development of this intravital imaging tool may be used to identify a mechanism for dendritic transmembrane protein trafficking, to observe the half-life of drugs or probes within the LCS, and to establish novel targets for modulating osteocyte mechanosensitivity to prevent or ameliorate bone metabolic disease states.

## Methods

### Animals

For the timelapse experiments, skeletally mature (16-20 weeks old) male and female wild-type mice on a C57BL/6 background were used (Jackson Laboratory and Charles River Laboratory). Male and female mouse experiments were performed as a single cohort when possible. Some groups were performed up to a month apart due to age differences, but all methodology remained constant. For the Ca^2+^ signaling with loading experiments, a mouse line with osteocyte-targeted expression of the genetically encoded Ca^2+^ indicator (GECI) GCaMP6f (Chen 2013) was used. These mice were bred in house by crossing Ai95-D mice [B6J.Cg-Gt(ROSA)26Sortm95.1(CAGGCaMP6f)Hze/MwarJ; JAX Labs] with DMP1/Cre mice [B6N.FVB-Tg(Dmp1-cre)1Jqfe/BwdJ; Jackson Labs], which have Cre recombinase driven by the DMP1 promoter, a gene predominantly expressed in osteocytes(59). Use of these mice for osteocyte Ca^2+^ signaling has been well established in our lab(33, 58). After experimental use, mice were euthanized via cervical dislocation while under isoflurane anesthesia. All methods and procedures were approved by the Institutional Animal Care and Use Committees of Cornell University. All methods and procedures are reported in accordance with ARRIVE guidelines and in accordance with relevant guidelines and regulations.

### Synthesis of Ultrasmall cRGD- and TAT-Functionalized Core–Shell Silica Nanoparticles

Ultrasmall fluorescent core–shell silica nanoparticles (C′ Dots) with covalently encapsulated Cy5 dye (derived from Cy5-maleimide; Lumiprobe) were synthesized in water as previously described(60, 61). Briefly, the Cy5 dye was conjugated to (3-mercaptopropyl)trimethoxysilane (MPTMS; Sigma– Aldrich), producing a net positively charged fluorophore that was incorporated into a silica core formed from tetramethyl orthosilicate (TMOS; Sigma–Aldrich). The particles were subsequently coated with a poly(ethylene glycol) (PEG) shell using methoxy-PEG(6–9)-silane (*∼* 500 g/mol; Gelest). Two varieties of C Dots were used in these experiments as described below. Integrin-targeting C′ Dots were surface functionalized with cyclic(arginine–glycine– aspartic acid–D-tyrosine–cysteine) peptide, c(RGDyC), to produce c(RGDyC)-PEG-Cy5-C′ Dots (hereafter referred to as RGD-C′ Dots). RGD-C Dots were functionalized by adding cRGDyC-PEG-silane to the particle core during the PEGylation step, as described in detail elsewhere(60, 62).

Membrane penetrating C Dots were surface functionalized with human immunodeficiency virus (HIV) TAT peptides (TAT_PEG-Cy5-C Dots) referred to as TAT-C Dots. TAT-C Dots are known to be capable of endosomal escape and were prepared similar to previously reported methods(63), but using an improved click chemistry approach. The TAT peptide sequence used was 5-azidopentanoyl – RKKRRQRRR-NH2 (Biosynth). The terminal azido group of the peptide was clicked onto the C’Dot surface functionalized with dibenzocyclooctyne (DBCO) via azide-alkyne cycloaddition. The DBCO functionalized C Dots were finalized using post-PEGylation surface modification by insertion (PPSMI) method as described in detail elsewhere(64). To that end, small functional aminopropyltrimethoxysilane (APTMS) was first inserted between PEGs on the silica surface. Resulting NH2-C’ dots were then reacted with DBCO-PEG4-N-hydroxysuccinimidyl ester (DBCO-PEG4-NHS ester, Santa Cruz Biotechnology) yielding DBCO-C Dots.

From particle characterization efforts employing a combination of fluorescence correlation spectroscopy (FCS) and UV-Vis absorbance data, with methods described elsewhere41, RGD- and TAT-C Dots exhibited hydrodynamic radii of 5.5, and 6.0nm, respectively, and number of dyes per C’Dot of 2.1, and 2.0, respectively. Furthermore, RGD-C Dots were estimated to have 20 cRGDyC ligands per particle and TAT-C Dots were estimated to have 10 TAT (5-azido-pentanoyl – RKKRRQRRR-NH2) peptides per particle.

### Endocytosis Modulating Small Molecule Compound

Water-soluble oligosaccharide methyl-β-cyclodextrin (MβCD, Sigma) binds to and removes cholesterol from the plasma membrane and is used to evaluate the impact of cholesterol depletion and cholesterol-mediated lipid raft-based endocytosis inhibition(34). M*β*CD was diluted from a powder to its working concentration of 30mM in 1x DPBS. M*β*CD was kept at 4and brought to room temperature before use. M*β*CD was used to disrupt plasma membrane cholesterol and endocytosis in the timelapse and Ca^2+^ signaling with loading experiments.

### MβCD Injection and Incubation

To inject M*β*CD, mice were anesthetized with 2-3% isoflurane mixed with medical grade air in an induction chamber for 3 minutes. Restraint of the mice was necessary to maintain reliable reproduction of the precise injection procedure. During injection, mice were kept under 2% isoflurane with a nose cone. A 10µL bolus of 30mM M*β*CD was subcutaneously injected above the third metatarsal (MT3) using a specialized low volume syringe (Hamilton) 30 minutes before additional injection or imaging. A local injection was used to minimize the off-target endocrine effects of a systemic injection. A 10µL bolus of 1x DPBS was used on control groups in the Ca^2+^ signaling with loading experiments (n=8-9/group). No control bolus was used in the timelapse experiments. Mice were allowed normal cage activity during the 30-minute pre-incubation time.

### Quantification of Membrane Cholesterol

A 15µL bolus of M*β*CD (30mM) or 1x DPBS control was injected subcutaneously above the MT3 of GCaMP6 mice as described in Section 4.4 and incubated for 30 minutes prior to sacrifice and MT3 dissection. MT3 bones were fixed overnight in Z-Fix, rinsed in PBS x3, then put in 70% ethanol at 4°C. Prior to cholesterol staining, MT3s were dehydrated in 30% sucrose overnight and then cryosectioned (6µm thickness)(65). Slides were rehydrated in 1xPBS for 12 minutes prior to staining. A fluorescent Filipin III stain was used to quantify cellular cholesterol levels(66), using an adapted protocol from Ma et al.(67). A 50µg/mL aliquot of Filipin III (Sigma Aldrich) was warmed to room temp from -20and then incubated with fixed MT3 cryosections for 30 minutes followed by a 3 minute 1xPBS rinse and cover slipping with Eukitt mounting media. Filipin III signal (DAPI filter) and GCaMP6 signal (GFP filter) were excited and collected using a fluorescent microscope (Keyence). Normalized fluorescence intensity of Filipin III signal in osteocytes was compared between control and M*β*CD treated groups for both sexes (ImageJ, NIH). GCaMP6 signal was used to identify osteocyte ROIs and max z-projections of 3 frames 1.25µm apart were analyzed for fluorescence for each group.

### Injection and Incubation of Fluorescent Nanoparticles and Membrane Markers

To inject the fluorescent markers for intravital imaging, mice were anesthetized as described anpve. For the timelapse experiments, M*β*CD treated or naïve mice were subcutaneously injected above the MT3 with 10µL of 10µM RGD- or TAT-C Dots and 10µL of the fluorescent cell membrane marker FM1-43 (ThermoFisher). C Dots and FM1-43 were incubated for 5 minutes and mice were kept under nose cone anesthesia before imaging (n=3/group/sex).

### Metatarsal Isolation Surgery

MT3 bones were isolated as previously described(31–33). Briefly, while mice were anesthetized, a shallow incision was made between the second and third metatarsal on the dorsal aspect of the mouse hind paw, and overlying tendons were removed. The MT3 was then functionally isolated from the rest of the paw with a stainless-steel pin beneath the mid-diaphysis of the bone, leaving primary vasculature at the proximal and distal palmar aspects intact. The bone was secured in a 3-point bending configuration and submerged a bath of 1X Phosphate Buffered Saline, Dulbecco’s Formula (DPBS, Thermo Scientific). Room temperature DPBS was used in the nanoparticle loading experiment and the Ca^2+^ signaling with loading experiment. For the timelapse study, a small immersion heater was used to maintain a DPBS bath temperature of 35throughout the experiment. This was done to support a higher rate of endocytosis and membrane mobility for timelapse imaging(68). Mice were continuously anesthetized with a nose cone at 1.5-2% isoflurane during all subsequent imaging.

### Metatarsal Loading

In the Ca^2+^ signaling with loading experiment, cyclic haversine loading at a previously calibrated strain of 1000µε was applied to the surgically isolated MT3(31, 69, 70). This strain level is within the range of strains that have been reported during physiological activities from*in vivo* strain gauge studies, with strains up to 2000µε characteristic of habitual activities(71, 72). A 15 minute wait time was used in between each loading session to account for the refractory period in osteocytes(73, 74). For each session, cyclic haversine loading was applied at 1Hz for 60 seconds. In the Ca^2+^ signaling with loading experiment, 8 sessions were used.

### Intravital Imaging

For all experiments, fluorescent signal inside the functionally isolated MT3 was visualized with multiphoton microscopy (MPM) (Bergamo II, Thorlabs) using a Ti-Sapphire laser (Coherent Chameleon Discovery NX) and 20x immersion objective (XLUMPLFLN, Olympus). Fluorescent excitation and imaging techniques were distinct for each experiment as described below.

### Timelapse Experiment

To observe the multiplexed signal of FM1-43 and Cy5 C Dots, 920nm wavelength excitation was used and emission was acquired with a 525±25nm bandpass filter (FM1-43) and a >647nm long pass filter (Cy5 C Dots). Images were captured at 1024×1024 pixel density and 0.292 pixel resolution. To track membrane and nanoparticle fluorescent signal variation and trafficking, images were taken at the same ROI 20µm below the bone surface every 60 seconds for 60 minutes at averaging of 15 frames per image. This imaging window includes the known timeframe for endocytic trafficking of 10-20 minutes.

### Calcium Signaling with Loading Experiment

To capture fluctuations in GCaMP6f Ca^2+^ signaling in osteocytes during physiological loading, 920nm excitation with a 525±25nm bandpass filter was used. Streaming images were captured at 512×512 pixel density, 0.292µm pixel resolution, frame averaging of 3 with a sampling rate of 29.4Hz, leading to final frame rate of 9.8 fps. Images were collected for 60 seconds prior to loading (non-loaded), 60 seconds during loading (loaded), and 30 seconds after loading (resting). This resulted in 1500 8-bit TIF images over 150 seconds for each loading bout. To correct for small deflections in the MT3 bone from loading, a motion correction system was utilized (PI E-709.SRG and ThorLabs Precision Controller)36.

### Image Analysis – Time Lapse Experiment

Variation in signal and trafficking activity was quantified in ImageJ (NIH). Analysis was repeated separately for both channels (FM1-43 membrane dye and C’Dot nanoparticle signal). Rectangular cell ROIs were identified prior to processing. Timelapse image cell ROI mean intensities were quantified across the full experiment using the ROI manager MultiMeasure tool. Variation in signal over time was assessed using the mean cell ROI intensity of a standard deviation z-projection that creates a single image by mapping the standard deviation in signal for each pixel across the full experiment. Each projection was calibrated to the same intensity range. Intensity analysis was normalized to a background intensity ROI for each channel and mouse. Qualitative analysis of membrane and nanoparticle trafficking was also performed. Overlayed channels were visualized through time and the number of cells with punctate signal (‘vesicle’) motion or signal closure over time was counted per mouse. Four scorers were blinded to the data groups and the average whole number value for each metric was used. If one scorer had an outlying yes/no response compared to the other three, their score for that metric was not counted.

### Image Analysis – Calcium Signaling with Loading Experiment

Intensity measurements were performed by postprocessing images using ImageJ (NIH). In-plane movements in the x and y were removed using the template matching plugin. ROIs were selected with a semi-automated process. A mask was chosen using thresholding selected by the user after projecting across the z with mean intensity followed by enhancing the contrast and applying a gaussian blur with sigma of 1. Size exclusion was performed, filtering ROIs for only 150-650 pixels. Finally, final adjustments to ROIs were made by eye, deleting and adding ROIs as needed that were not properly captured by thresholding. An ROI indicating the background autofluorescence was included in an area where no cells were present. The average pixel intensity for each ROI was collected across all 1500 time series frames and output as a csv.

In MATLAB, ROIs that exceeded a cutoff of the mean background intensity plus 3 standard deviations were included in the final analysis as fluorescent osteocytes. The intensities for each cell of interest were detrended linearly to correct for photobleaching and normalized to the mean intensity for that cell over a 40 second period prior to loading. Peaks and troughs were identified using islocalmax and islocalmin in MATLAB with a prominence of 0.01. Amplitudes across the non-loaded and loaded intervals were computed by subtracting coupled troughs from peaks. Changes in amplitude fluorescent intensity were computed as follows: 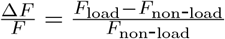 indicates the amplitude during loading and 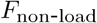 indicates the amplitude without loading. Responding cells were defined as cells showing a > 25% increase in Δ*F/F* .

### Statistical Analyses

Comparisons in the cholesterol Filipin III stain experiment were made with a one-way ANOVA with multiple comparisons and Sidak’s correction. For the timelapse experiment, one-way ANOVA’s were also used to compare channels and treated vs non-treated groups. For the Ca^2+^ signaling with loading, linear regressions were used to compare the M*β*CD and control groups across the repeated loading sessions. Statistical significance was determined as p<0.05. Analyses were performed in GraphPad Prism version 9.0 for Mac OS X (GraphPad Software).

## ACKNOWLEDGEMENTS

We would like to thank the Cornell CARE facility. Work supported by NIH NIAMS R01AR087805-01

## Supplementary Figures

**Fig. S1.**
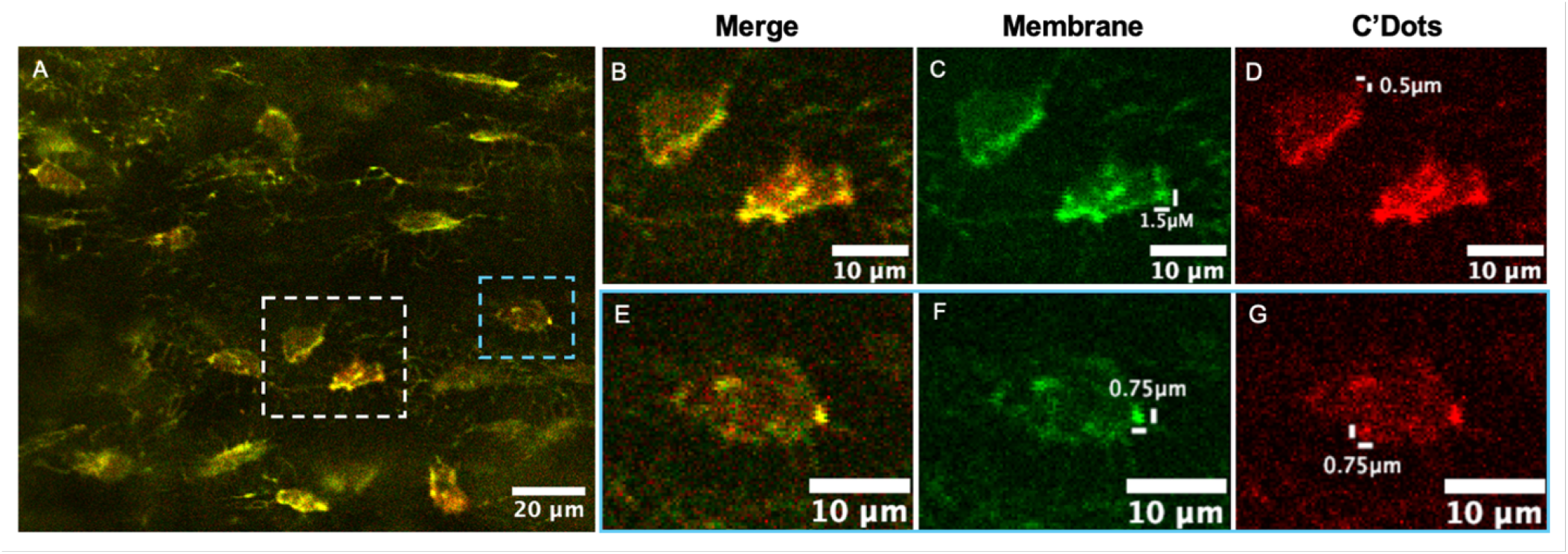
Multiplexing of exogenous fluorescent signal delivered by local subcutaneous injection enables rapid visualization of osteocyte membrane dynamics and nanoparticle uptake and trafficking. Representative merged image of membrane (green) and RGD-C^2+^ Dot nanoparticle signal in MT3 cortical bone osteocytes *in vivo* following a 5 min incubation (A). Scale bar = 20 *µ*m. Cells exhibiting submicron-resolution punctate signal are indicated by dashed boxes and shown at higher magnification with merged and separated fluorescence channels (white box, B–D; blue box, E–F). Scale bars = 10 *µ*m.

**Fig. S2.**
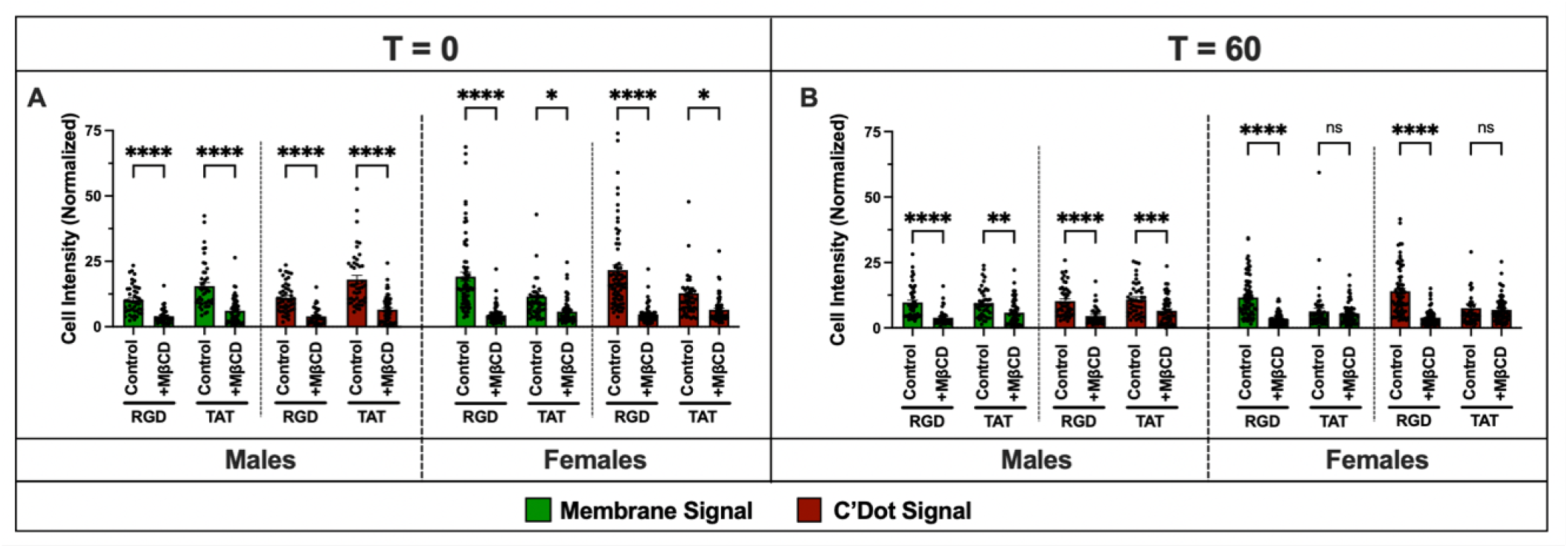
The effects of M*β*CD treatment on initial (0 min, A) and final (60 min, B) cellular fluorescence intensity. Membrane (green) and C^2+^ Dot (red) fluorescence intensities were compared across C^2+^ Dot functionalizations and sexes using one-way ANOVAs with multiple comparisons. *n* = 3 mice per group, with 10–30 cells analyzed per mouse. Individual data points represent measurements from single cells. Error bars represent SEM. \**p <* 0.05, \*\**p <* 0.01, \*\*\**p <* 0.001, \*\*\*\**p <* 0.0001.

**Fig. S3.**
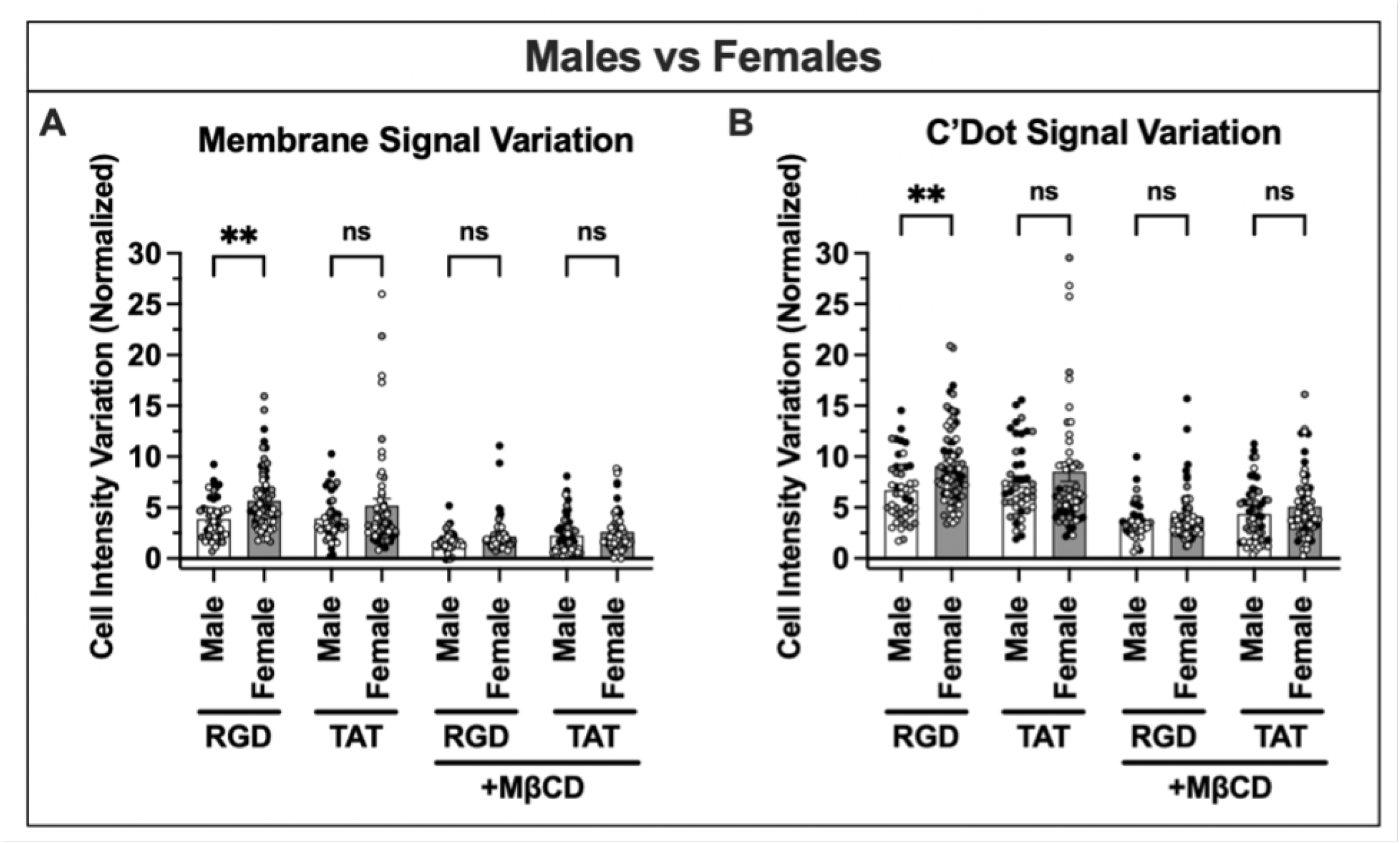
The effects of sex on signal intensity variation for membrane (A) and C Dot (B) signals. Male and female mice were compared within control and M*β*CD-treated groups using one-way ANOVAs with multiple comparisons. *n* = 3 mice per group, with 10–30 cells analyzed per mouse. Individual data points represent measurements from single cells, with colors indicating individual mice. Error bars represent SEM. \**p <* 0.05, \*\**p <* 0.01, \*\*\**p <* 0.001, \*\*\*\**p <* 0.0001.

## REFERENCE

1. Clinton T Rubin and Lance E Lanyon. Regulation of bone formation by applied dynamic loads. The Journal of Bone Joint Surgery, 66(3):397–402, 1984. This paper shows that changes in strain distribution, independent of rate and magnitude, leads to periosteal bone formation. Suggests that bone cells only require a short bout of loading to maintain bone mass.

2. Charles H Turner. Three rules for bone adaptation to mechanical stimuli. Bone, 23(5):399–407, 1998.

3. Sheldon Weinbaum, Stephen C Cowin, and Yu Zeng. A model for the excitation of osteocytes by mechanical loading-induced bone fluid shear stresses. Journal of Biomechanics, 27(3):339–360, 1994.

4. Charles H Turner, Mark R Forwood, Jae-Young Rho, and Tomoaki Yoshikawa. Mechanical loading thresholds for lamellar and woven bone formation. Journal of Bone and Mineral Research, 9(1):87–97, 1994.

5. Lynda F Bonewald. The amazing osteocyte. Journal of Bone and Mineral Research, 26 (2):229–238, 2011. doi: 10.1002/jbmr.320. Good original finding references. Good listing of actin binding protein references. Great references for current and older osteocyte information. Use when writing!!

6. Christopher Price, Xiaozhou Zhou, Wen Li, and Liyun Wang. Real-time measurement of solute transport within the lacunar-canalicular system of mechanically loaded bone: direct evidence for load-induced fluid flow. Journal of Bone and Mineral …, 26(2):277–285, 2011. doi: 10.1002/jbmr.211.

7. Mia M Thi, Sylvia O Suadicani, Mitchell B Schaffler, Sheldon Weinbaum, and David C Spray. Mechanosensory responses of osteocytes to physiological forces occur along processes and not cell body and require αvβ3 integrin. Proceedings of the National Academy of Sciences of the United States of America, 110(52):21012–21017, 12 2013. doi: 10.1073/pnas.1321210110.

8. Stefaan W Verbruggen, Ted J Vaughan, and Laoise M McNamara. Fluid flow in the osteocyte mechanical environment: a fluid–structure interaction approach. Biomechanics and Modeling in Mechanobiology, 13(1):85–97, 2013. doi: 10.1007/s10237-013-0487-y.

9. Yuefeng Han, Stephen C Cowin, Mitchell B Schaffler, and Sheldon Weinbaum.Mechanotransduction and strain amplification in osteocyte cell processes. Proceedings of the National Academy of Sciences, 101(47):16689–16694, 11 2004.

10. S. J. Singer. A fluid lipid-globular protein mosaic model of membrane structure*. Annals of the New York Academy of Sciences, 195(1):16–23, 1972. ISSN 0077-8923. doi: 10.1111/j.1749-6632.1972.tb54780.x.

11. Garth L. Nicolson. The fluid—mosaic model of membrane structure: Still relevant to understanding the structure, function and dynamics of biological membranes after more than 40years. Biochimica et Biophysica Acta (BBA) - Biomembranes, 1838(6):1451–1466, 2014. ISSN 0005-2736. doi: 10.1016/j.bbamem.2013.10.019.

12. Ken Jacobson, Erin D. Sheets, and Rudolf Simson. Revisiting the fluid mosaic model of membranes. Science, 268(5216):1441–1442, 1995. ISSN 0036-8075. doi: 10.1126/science.7770769.

13. Anabel-Lise Le Roux, Xarxa Quiroga, Nikhil Walani, Marino Arroyo, and Pere Roca-Cusachs. The plasma membrane as a mechanochemical transducer. Philosophical Transactions of the Royal Society B, 374(1779):20180221, 2019. ISSN 0962-8436. doi: 10.1098/rstb.2018.0221.

14. Rebecca Yarwood, John Hellicar, Philip G. Woodman, and Martin Lowe. Membrane trafficking in health and disease. Disease Models Mechanisms, 13(4):dmm043448, 2020. ISSN 1754-8403. doi: 10.1242/dmm.043448.

15. Upasana Mukhopadhyay, Tithi Mandal, Madhura Chakraborty, and Bidisha Sinha. The plasma membrane and mechanoregulation in cells. ACS Omega, 9(20):21780–21797, 2024. ISSN 2470-1343. doi: 10.1021/acsomega.4c01962.

16. Anqi Zhou, Bao Xue, Jiayi Zhong, Junjun Liu, Ran Peng, Fan Wang, Yuan Zhou, Jielin Tang, Qi Yang, and Xinwen Chen. Cholesterol-rich lipid rafts mediate endocytosis as a common pathway for respiratory syncytial virus entry into different host cells. Microbiology Spectrum, 13(9):e01192–25, 2025. ISSN 2165-0497. doi: 10.1128/spectrum.01192-25.

17. Nyein Nyein Chan, Manabu Yamazaki, Satoshi Maruyama, Tatsuya Abé, Kenta Haga, Masami Kawaharada, Kenji Izumi, Tadaharu Kobayashi, and Jun-ichi Tanuma. Cholesterol is a regulator of cav1 localization and cell migration in oral squamous cell carcinoma. International Journal of Molecular Sciences, 24(7):6035, 2023. ISSN 1422-0067. doi: 10.3390/ijms24076035.

18. Philippe G. Frank, Yves L. Marcel, Margery A. Connelly, Douglas M. Lublin, Vivian Franklin, David L. Williams, and Michael P. Lisanti. Stabilization of caveolin-1 by cellular cholesterol and scavenger receptor class b type i. Biochemistry, 41(39):11931–11940, 2002. ISSN 0006-2960. doi: 10.1021/bi0257078.

19. Arikta Biswas, Purba Kashyap, Sanchari Datta, Titas Sengupta, and Bidisha Sinha. Cholesterol depletion by mβcd enhances cell membrane tension and its variations-reducing integrity. Biophysical Journal, 116(8):1456–1468, 2019. ISSN 0006-3495. doi: 10.1016/j.bpj.2019.03.016.

20. Raphael Zidovetzki and Irena Levitan. Use of cyclodextrins to manipulate plasma membrane cholesterol content: evidence, misconceptions and control strategies. Biochimica et biophysica acta, 1768(6):1311–1324, 2007. ISSN 0006-3002. doi: 10.1016/j.bbamem.2007.03.026.

21. Don Armstrong and Raphael Zidovetzki. Amplification of diacylglycerol activation of protein kinase c by cholesterol. Biophysical Journal, 94(12):4700–4710, 2008. ISSN 0006-3495. doi: 10.1529/biophysj.107.121426.

22. Subburaj ILANGUMARAN and Daniel C. HOESSLI. Effects of cholesterol depletion by cyclodextrin on the sphingolipid microdomains of the plasma membrane. Biochemical Journal, 335(2):433–440, 1998. ISSN 0264-6021. doi: 10.1042/bj3350433.

23. Mohammad Niroobakhsh, Yixia Xie, Sarah L. Dallas, Mark L. Johnson, and Thiagarajan Ganesh. Analysis and imaging of osteocytes. Journal of visualized experiments : JoVE, 213:10.3791/64699, 2024. ISSN 1940-087X. doi: 10.3791/64699.

24. T. J. Vaughan, M. G. Haugh, and L. M. McNamara. A fluid–structure interaction model to characterize bone cell stimulation in parallel-plate flow chamber systems. Journal of The Royal Society Interface, 10(81):20120900, 2013. ISSN 1742-5689. doi: 10.1098/rsif.2012.0900.

25. Luoding () Zhu, Jared Barber, Robert Zigon, Sungsoo () Na, and Hiroki () Yokota. Modeling and simulation of interstitial fluid flow around an osteocyte in a lacuno-canalicular network. Physics of Fluids, 34(4):041906, 2022. ISSN 1070-6631. doi: 10.1063/5.0085299.

26. Yan Li, Chunxi Ge, Jason P Long, Dana L Begun, Jose A Rodriguez, Steven A Goldstein, and Renny T Franceschi. Biomechanical stimulation of osteoblast gene expression requires phosphorylation of the runx2 transcription factor. Journal of Bone and Mineral Research, 27(6):1263–1274, 05 2012. doi: 10.1002/jbmr.1574.

27. Robin Michael Delaine-Smith, Behzad Javaheri, Jennifer Helen Edwards, Marisol Vazquez, and Robin Mark Howard Rumney. Preclinical models for in vitro mechanical loading of bone-derived cells. BoneKEy Reports, 4:728, 2015. ISSN 2047-6396. doi: 10.1038/bonekey.2015.97.

28. Taiji Adachi, Yuki Aonuma, Shin-ichi Ito, Mototsugu Tanaka, Masaki Hojo, Teruko Takano-Yamamoto, and Hiroshi Kamioka. Osteocyte calcium signaling response to bone matrix deformation. Journal of Biomechanics, 42(15):2507–2512, 11 2009. doi: 10.1016/j.jbiomech.2009.07.006.

29. Da Jing, Andrew D Baik, X Lucas Lu, Bin Zhou, Xiaohan Lai, Liyun Wang, Erping Luo, and X Edward Guo. In situ intracellular calcium oscillations in osteocytes in intact mouse long bones under dynamic mechanical loading. FASEB journal : official publication of the Federation of American Societies for Experimental Biology, 28(4):1582–1592, 2014. doi: 10.1096/fj.13-237578.

30. Andrea E Morrell, Genevieve N Brown, Samuel T Robinson, Rachel L Sattler, Andrew D Baik, Gehua Zhen, Xu Cao, Lynda F Bonewald, Weiyang Jin, Lance C Kam, and X Edward Guo. Mechanically induced ca2+ oscillations in osteocytes release extracellular vesicles and enhance bone formation. Bone Research, 6(1):1–11, 03 2018. doi: 10.1038/s41413-018-0007-x.

31. Karl J Lewis, Dorra Frikha-Benayed, Joyce Louie, Samuel Stephen, David C Spray, Mia M Thi, Zeynep Seref-Ferlengez, Robert J Majeska, Sheldon Weinbaum, and Mitchell B Schaffler. Osteocyte calcium signals encode strain magnitude and loading frequency in vivo. Proceedings of the National Academy of Sciences of the United States of America, 114 (44):11775–11780, 10 2017. doi: 10.1073/pnas.1707863114.

32. Melia D. Matthews, Emily Cook, Nada Naguib, Ulrich B. Wiesner, and Karl J. Lewis. Intravital imaging of osteocyte integrin dynamics with locally injectable fluorescent nanoparticles. Bone, 174:116830, 2023. ISSN 8756-3282. doi: 10.1016/j.bone.2023.116830.

33. Melia D. Matthews, Alexander Saffari, Nuzhat Mukul, Lanlan Hai, Nada Naguib, Ulrich B. Wiesner, and Karl J. Lewis. Real-time visualization and modulation of endocytic dynamics in osteocytes in vivo. Scientific Reports, 15(1):31454, 2025. ISSN 2045-2322. doi: 10.1038/s41598-025-05735-1.

34. Saleemulla Mahammad and Ingela Parmryd. Cholesterol depletion using methyl-β-cyclodextrin. In [“Owen and Dylan M.”], editors, Methods in Membrane Lipids, volume 1232 of Methods in molecular biology (Clifton, N.J.), pages 91–102. Springer, New York, NY, 2014. ISBN 9781493917518. doi: 10.1007/978-1-4939-1752-5_8.

35. Hatem Tallima and Rashika El Ridi. Methyl-β-cyclodextrin treatment and filipin staining reveal the role of cholesterol in surface membrane antigen sequestration of schistosoma mansoni and s. haematobium lung-stage larvae. Journal of Parasitology, 91(3):720–725, 2005. ISSN 0022-3395. doi: 10.1645/ge-439r.

36. Mitsuhiro Abe and Toshihide Kobayashi. Imaging cholesterol depletion at the plasma membrane by methyl-β-cyclodextrin. Journal of Lipid Research, 62, 2021. ISSN 0022-2275. doi: 10.1016/j.jlr.2021.100077.

37. Pietro Ridone, Elvis Pandzic, Massimo Vassalli, Charles D. Cox, Alexander Macmillan, Philip A. Gottlieb, and Boris Martinac. Disruption of membrane cholesterol organization impairs the activity of piezo1 channel clusters. Journal of General Physiology, 152(8): e201912515, 2020. ISSN 0022-1295. doi: 10.1085/jgp.201912515.

38. Stacy A. Francis, Joan M. Kelly, Joanne Mccormack, Rick A. Rogers, Jean Lai, Eveline E. Schneeberger, and Robert D. Lynch. Rapid reduction of mdck cell cholesterol by methyl-β-cyclodextrin alters steady state transepithelial electrical resistance. European Journal of Cell Biology, 78(7):473–484, 1999. ISSN 0171-9335. doi: 10.1016/s0171-9335(99)80074-0.

39. Siv Kjersti Rodal, Grethe Skretting, Øystein Garred, Frederik Vilhardt, Bo van Deurs, and Kirsten Sandvig. Extraction of cholesterol with methyl-β-cyclodextrin perturbs formation of clathrin-coated endocytic vesicles. Molecular Biology of the Cell, 10(4):961–974, 1999. ISSN 1059-1524. doi: 10.1091/mbc.10.4.961.

40. Dries Vercauteren, Roosmarijn E. Vandenbroucke, Arwyn T. Jones, Joanna Rejman, Joseph Demeester, Stefaan C. De Smedt, Niek N. Sanders, and Kevin Braeckmans. The use of inhibitors to study endocytic pathways of gene carriers: Optimization and pitfalls. Molecular Therapy, 18(3):561–569, 2010. ISSN 1525-0016. doi: 10.1038/mt.2009.281.

41. Min Cheng, Rui Zhang, Jianshu Li, Wenyuan Ma, Linrun Li, Na Jiang, Bingxin Liu, Jing Wu, Nan Zheng, and Zhiwei Wu. Mβcd inhibits sftsv entry by disrupting lipid raft structure of the host cells. Antiviral Research, 231:106004, 2024. ISSN 0166-3542. doi: 10.1016/j.antiviral.2024.106004.

42. Yanghui Xing, Yan Gu, Li-Chong Xu, Christopher A. Siedlecki, Henry J. Donahue, and Jun You. Effects of membrane cholesterol depletion and gpi-anchored protein reduction on osteoblastic mechanostransduction. Journal of Cellular Physiology, 226(9):2350–2359, 2011. ISSN 0021-9541. doi: 10.1002/jcp.22579.

43. Monika Lakk, Grace F. Hoffmann, Aruna Gorusupudi, Eric Enyong, Amy Lin, Paul S. Bernstein, Trine Toft-Bertelsen, Nanna MacAulay, Michael H. Elliott, and David Križaj. Membrane cholesterol regulates trpv4 function, cytoskeletal expression, and the cellular response to tension. Journal of Lipid Research, 62:100145, 2021. ISSN 0022-2275. doi: 10.1016/j.jlr.2021.100145.

44. Laoise M McNamara, Robert J Majeska, Sheldon Weinbaum, Victor Friedrich, and Mitchell B Schaffler. Attachment of osteocyte cell processes to the bone matrix. The Anatomical Record: Advances in Integrative Anatomy and Evolutionary Biology, 292(3): 355–363, 2009. doi: 10.1002/ar.20869.

45. Pamela Cabahug-Zuckerman, Randy F Stout Jr., Robert J Majeska, Mia M Thi, David C Spray, Sheldon Weinbaum, and Mitchell B Schaffler. Potential role for a specialized β 3integrin-based structure on osteocyte processes in bone mechanosensation. Journal of orthopaedic research, 118:733–11, 11 2017. doi: 10.1002/jor.23792.

46. Paulina Moreno-Layseca, Jaroslav Icha, Hellyeh Hamidi, and Johanna Ivaska. Integrin trafficking in cells and tissues. Nature Cell Biology, 21(2):122–132, 2019. ISSN 1465-7392. doi: 10.1038/s41556-018-0223-z.

47. Viola Hélène Lobert, Andreas Brech, Nina Marie Pedersen, Jørgen Wesche, Angela Oppelt, Lene Malerød, and Harald Stenmark. Ubiquitination of α5β1 integrin controls fibroblast migration through lysosomal degradation of fibronectin-integrin complexes. Developmental Cell, 19(1):148–159, 2010. ISSN 1534-5807. doi: 10.1016/j.devcel.2010.06.010.

48. Antti Arjonen, Jonna Alanko, Stefan Veltel, and Johanna Ivaska. Distinct recycling of active and inactive β1 integrins. Traffic, 13(4):610–625, 2012. ISSN 1398-9219. doi: 10.1111/j.1600-0854.2012.01327.x.

49. Kayo Tanaka-Kamioka, Hiroshi Kamioka, Hans Ris, and Soo-Siang Lim. Osteocyte shape is dependent on actin filaments and osteocyte processes are unique actin-rich projections. Journal of Bone and Mineral Research, 13(10):1555–1568, 1998. ISSN 0884-0431. doi: 10.1359/jbmr.1998.13.10.1555.

50. Li-Dan You, Sheldon Weinbaum, Stephen C Cowin, and Mitchell B Schaffler. Ultrastructure of the osteocyte process and its pericellular matrix. The Anatomical Record: Advances in Integrative Anatomy and Evolutionary Biology, 278A(2):505–513, 2004. doi: 10.1002/ar.a.20050.

51. Rosa M. Guerra, Velia M. Fowler, and Liyun Wang. Osteocyte dendrites: How do they grow, mature, and degenerate in mineralized bone? Cytoskeleton, 82(9):556–570, 2025. ISSN 1949-3592. doi: 10.1002/cm.21964.

52. Shigeo Takamori, Matthew Holt, Katinka Stenius, Edward A. Lemke, Mads Grønborg, Dietmar Riedel, Henning Urlaub, Stephan Schenck, Britta Brügger, Philippe Ringler, Shirley A. Müller, Burkhard Rammner, Frauke Gräter, Jochen S. Hub, Bert L. De Groot, Gottfried Mieskes, Yoshinori Moriyama, Jürgen Klingauf, Helmut Grubmüller, John Heuser, Felix Wieland, and Reinhard Jahn. Molecular anatomy of a trafficking organelle. Cell, 127(4): 831–846, 2006. ISSN 0092-8674. doi: 10.1016/j.cell.2006.10.030.

53. Gabrielle Larocque and Stephen J. Royle. Integrating intracellular nanovesicles into integrin trafficking pathways and beyond. Cellular and Molecular Life Sciences, 79(6):335, 2022. ISSN 1420-9071. doi: 10.1007/s00018-022-04371-6.

54. Sakhr A. Murshid, Hiroshi Kamioka, Yoshihito Ishihara, Ryoko Ando, Yasuyo Sugawara, and Teruko Takano-Yamamoto. Actin and microtubule cytoskeletons of the processes of 3d-cultured mc3t3-e1 cells and osteocytes. Journal of Bone and Mineral Metabolism, 25(3): 151–158, 2007. ISSN 1435-5604. doi: 10.1007/s00774-006-0745-5.

55. Hai Qing, Laleh Ardeshirpour, Paola Divieti Pajevic, Vladimir Dusevich, Katharina Jähn, Shigeaki Kato, John Wysolmerski, and Lynda F Bonewald. Demonstration of osteocytic perilacunar/canalicular remodeling in mice during lactation. Journal of Bone and Mineral Research, 27(5):1018–1029, 2012. ISSN 0884-0431. doi: 10.1002/jbmr.1567.

56. Michael Sieverts, Cristal Yee, Minali Nemani, Dilworth Y. Parkinson, Tamara Alliston, and Claire Acevedo. Spatial control of perilacunar canalicular remodeling during lactation. Scientific Reports, 14(1):14655, 2024. ISSN 2045-2322. doi: 10.1038/s41598-024-63645-0.

57. John J Wysolmerski. Osteocytes remove and replace perilacunar mineral during reproductive cycles. Bone, 54(2):230–236, 2013. ISSN 8756-3282. doi: 10.1016/j.bone.2013.01.025.

58. Macy Mora-Antoinette, Andrea Garcia-Ortiz, Mariam Obaji, Alexander Saffari, Melia D. Matthews, Murtaza Wasi, and Karl J. Lewis. Cholinergic regulation of osteocyte mechanobiology: A paradigm for bone adaptation. Science Advances, 11(34):eads9720, 2025. ISSN 2375-2548. doi: 10.1126/sciadv.ads9720.

59. Yongbo Lu, Yixia Xie, Shubin Zhang, Vladimir Dusevich, Lynda F Bonewald, and Jian Q Feng. Dmp1-targeted cre expression in odontoblasts and osteocytes. Journal of Dental Research, 86(4):320–325, 04 2007. doi: 10.1177/154405910708600404.

60. Kai Ma, Carlie Mendoza, Margaret Hanson, Ulrike Werner-Zwanziger, Josef Zwanziger, and Ulrich Wiesner. Control of ultrasmall sub-10 nm ligand-functionalized fluorescent core–shell silica nanoparticle growth in water. Chemistry of Materials, 27(11):4119–4133, 2015. ISSN 0897-4756. doi: 10.1021/acs.chemmater.5b01222.

61. Kai Ma, Duhan Zhang, Ying Cong, and Ulrich Wiesner. Elucidating the mechanism of silica nanoparticle pegylation processes using fluorescence correlation spectroscopies. Chemistry of Materials, 28(5):1537–1545, 2016. ISSN 0897-4756. doi: 10.1021/acs.chemmater.6b00030.

62. Feng Chen, Kai Ma, Miriam Benezra, Li Zhang, Sarah M. Cheal, Evan Phillips, Barney Yoo, Mohan Pauliah, Michael Overholtzer, Pat Zanzonico, Sonia Sequeira, Mithat Gonen, Thomas Quinn, Ulrich Wiesner, and Michelle S. Bradbury. Cancer-targeting ultrasmall silica nanoparticles for clinical translation: Physicochemical structure and biological property correlations. Chemistry of Materials, 29(20):8766–8779, 2017. ISSN 0897-4756. doi: 10.1021/acs.chemmater.7b03033.

63. Jinhyang Choi, Andrew A. Burns, Rebecca M. Williams, Zongxiang Zhou, Andrea Flesken-Nikitin, Warren R. Zipfel, Ulrich Wiesner, and Alexander Y. Nikitin. Core-shell silica nanoparticles as fluorescent labels for nanomedicine. Journal of Biomedical Optics, 12(6):064007–064007–11, 2007. ISSN 1083-3668. doi: 10.1117/1.2823149.

64. Kai Ma and Ulrich Wiesner. Modular and orthogonal post-pegylation surface modifications by insertion enabling penta-functional ultrasmall organic-silica hybrid nanoparticles. Chemistry of Materials, 29(16):6840–6855, 2017. ISSN 0897-4756. doi: 10.1021/acs.chemmater.7b02009.

65. Huiliang Yang, Liang Wang, Kathleen Turajane, Lijun Wang, and Wentian Yang. A method for colocalizing lineage tracing reporter and rnascope signals on skeletal tissue section. RNA, 27(3):359–365, 2021. ISSN 1355-8382. doi: 10.1261/rna.077958.120.

66. Francesco Bruno, Serena Camuso, Elisabetta Capuozzo, and Sonia Canterini. The anti-fungal antibiotic filipin as a diagnostic tool of cholesterol alterations in lysosomal storage diseases and neurodegenerative disorders. Antibiotics, 12(1):122, 2023. ISSN 2079-6382. doi: 10.3390/antibiotics12010122.

67. Yiyang Ma, Yidan Pang, Chenglong Liu, Yuchen Tian, Kaiwen Zheng, Meng Yao, Xiaofeng Liu, Ruomu Cao, Yiwei Zhao, Zhikai Zheng, Weitao Jia, Daoyu Zhu, Hao Peng, Dajiang Du, Xinhua Qu, Chuan-ju Liu, Pei Yang, Yigang Huang, Changqing Zhang, and Junjie Gao. Mitochondria relay cholesterol signal exacerbates osteoarthritis in mice. Nature Communications, 16(1):10123, 2025. ISSN 2041-1723. doi: 10.1038/s41467-025-65689-w.

68. P. H. Weigel and J. A. Oka. Temperature dependence of endocytosis mediated by the asialoglycoprotein receptor in isolated rat hepatocytes. evidence for two potentially rate-limiting steps. The Journal of Biological Chemistry, 256(6):2615–2617, 1981. ISSN 0021-9258.

69. Karl J. Lewis, Pamela Cabahug-Zuckerman, James F. Boorman-Padgett, Jelena Basta-Pljakic, Joyce Louie, Samuel Stephen, David C. Spray, Mia M. Thi, Zeynep Seref-Ferlengez, Robert J. Majeska, Sheldon Weinbaum, and Mitchell B. Schaffler. Estrogen depletion on in vivo osteocyte calcium signaling responses to mechanical loading. Bone, 152:116072, 2021. ISSN 8756-3282. doi: 10.1016/j.bone.2021.116072.

70. Karl J Lewis, James F Boorman-Padgett, Macy Castaneda, David C Spray, Mia M Thi, and Mitchell B Schaffler. A fluorescent intravital imaging approach to study load-induced calcium signaling dynamics in mouse osteocytes. Journal of Visualized Experiments, (192), 2023. doi: 10.3791/64366.

71. D. B. Burr, C. Milgrom, D. Fyhrie, M. Forwood, M. Nyska, A. Finestone, S. Hoshaw, E. Saiag, and A. Simkin. In vivo measurement of human tibial strains during vigorous activity. Bone, 18(5):405–410, 1996. ISSN 8756-3282. doi: 10.1016/8756-3282(96)00028-2.

72. Clinton T Rubin. Skeletal strain and the functional significance of bone architecture. Calcified tissue international, 36(1):S11–S18, 1984.

73. Christopher R Jacobs, Clare E Yellowley,R R Davis, Zhiyi Zhou, John M Cimbala, and Henry J Donahue. Differential effect of steady versus oscillating flow on bone cells. Journal of Biomechanics, 31(11):969–976, 1998. doi: 10.1016/s0021-9290(98)00114-6.

74. Seth W. Donahue, Henry J. Donahue, and Christopher R. Jacobs. Osteoblastic cells have refractory periods for fluid-flow-induced intracellular calcium oscillations for short bouts of flow and display multiple low-magnitude oscillations during long-term flow. Journal of Biomechanics, 36(1):35–43, 2003. ISSN 0021-9290. doi: 10.1016/s0021-9290(02)00318-4.

